# Altered oligodendroglia and astroglia in chronic traumatic encephalopathy

**DOI:** 10.1101/2020.05.13.089086

**Authors:** K. Blake Chancellor, Sarah E. Chancellor, Joseph E. Duke-Cohan, Bertrand R. Huber, Thor D. Stein, Victor E. Alvarez, Benjamin W. Okaty, Susan M. Dymecki, Ann C. McKee

## Abstract

Chronic traumatic encephalopathy (CTE) is a progressive tauopathy found in contact sport athletes, military veterans, and others exposed to repetitive head impacts (RHI)^1–6^. White matter atrophy and axonal loss have been reported in CTE but have not been characterized on a molecular or cellular level^2,7,8^. Here, we present RNA sequencing profiles of cell nuclei from postmortem dorsolateral frontal white matter from eight individuals with neuropathologically confirmed CTE and eight age- and sex-matched controls. Analyzing these profiles using unbiased clustering approaches, we identified eighteen transcriptomically distinct cell groups (clusters), reflecting cell types and/or cell states, of which a subset showed differences between CTE and control tissue. Independent in situ methods applied on tissue sections adjacent to that used in the single-nucleus RNA-seq work yielded similar findings. Oligodendrocytes were found to be most severely affected in the CTE white matter samples; they were diminished in number and altered in relative proportions across subtype clusters. Further, the CTE-enriched oligodendrocyte population showed greater abundance of transcripts relevant to iron metabolism and cellular stress response. CTE tissue also demonstrated excessive iron accumulation histologically. Astrocyte alterations were more nuanced; total astrocyte number was indistinguishable between CTE and control samples, but transcripts associated with neuroinflammation were elevated in the CTE astrocyte groups as compared to controls. These results demonstrate specific molecular and cellular differences in CTE oligodendrocytes and astrocytes and may provide a starting point for the development of diagnostics and therapeutic interventions.

## INTRODUCTION

Chronic traumatic encephalopathy (CTE) is a progressive tauopathy associated with exposure to repetitive head impacts (RHI)^1–6^. CTE has been reported in contact sport athletes, military veterans and others exposed to neurotrauma^1,2,6^. The symptoms of CTE include mood and behavior abnormalities, cognitive impairment, and dementia^2,9–11^; however, like most neurodegenerative diseases, the diagnosis of CTE can only be made on post-mortem neuropathological evaluation, and there are no treatments. The distinctive hyperphosphorylated tau (p-tau) pathology of CTE primarily involves the frontal and temporal cortices^1,2,3^, although there are also profound changes in the white matter, including white matter rarefaction, blood brain barrier leakage, myelin and axonal loss, hemosiderin-laden macrophages and inflammation^2,6–8,12^.

We explored cell-type-specific mRNA profiles in white matter from subjects with neuropathologically-verified CTE compared to age-, sex-, and post-mortem interval (PMI)-matched controls by applying the method of single-nucleus RNA sequencing (snRNA-seq)^13–17^, in combination with complementary in situ histology, immunohistochemistry (IHC), immunofluorescence (IF), and single-molecule fluorescent mRNA in situ hybridization (smFISH) validation techniques. Our transcriptomic analyses yielded 24,735 individual brain nuclei RNA profiles. Our findings–the first at single-nucleus resolution in CTE–indicate differences in cell types, subpopulations, and gene expression in CTE white matter compared to controls.

## RESULTS

### Single-nucleus RNA sequencing of human postmortem CTE white matter

We performed molecular profiling of individual cell nuclei from fresh-frozen white matter tissue from the dorsolateral frontal cortex (Brodmann area (BA) 8/9). We collected nuclei from 16 age- and PMI-matched male individuals (Fig. 1a, Extended Data Fig. 1a-d). Clinical data and demographics are provided in Extended Data Table 1. Brain tissue was collected from 8 individuals diagnosed with CTE Stage II or III^2,3^ and 8 individuals with no neuropathological disease or known history of RHI. All 8 CTE subjects were former American football players (Extended Data Table 1) with no co-morbid neurodegenerative disease. CTE Stage II and III were selected for study, rather than early- or late-stage, reasoning that moderate disease would likely reveal transcriptomic changes most directly pertinent to the pathology of CTE.

**Figure 1.**
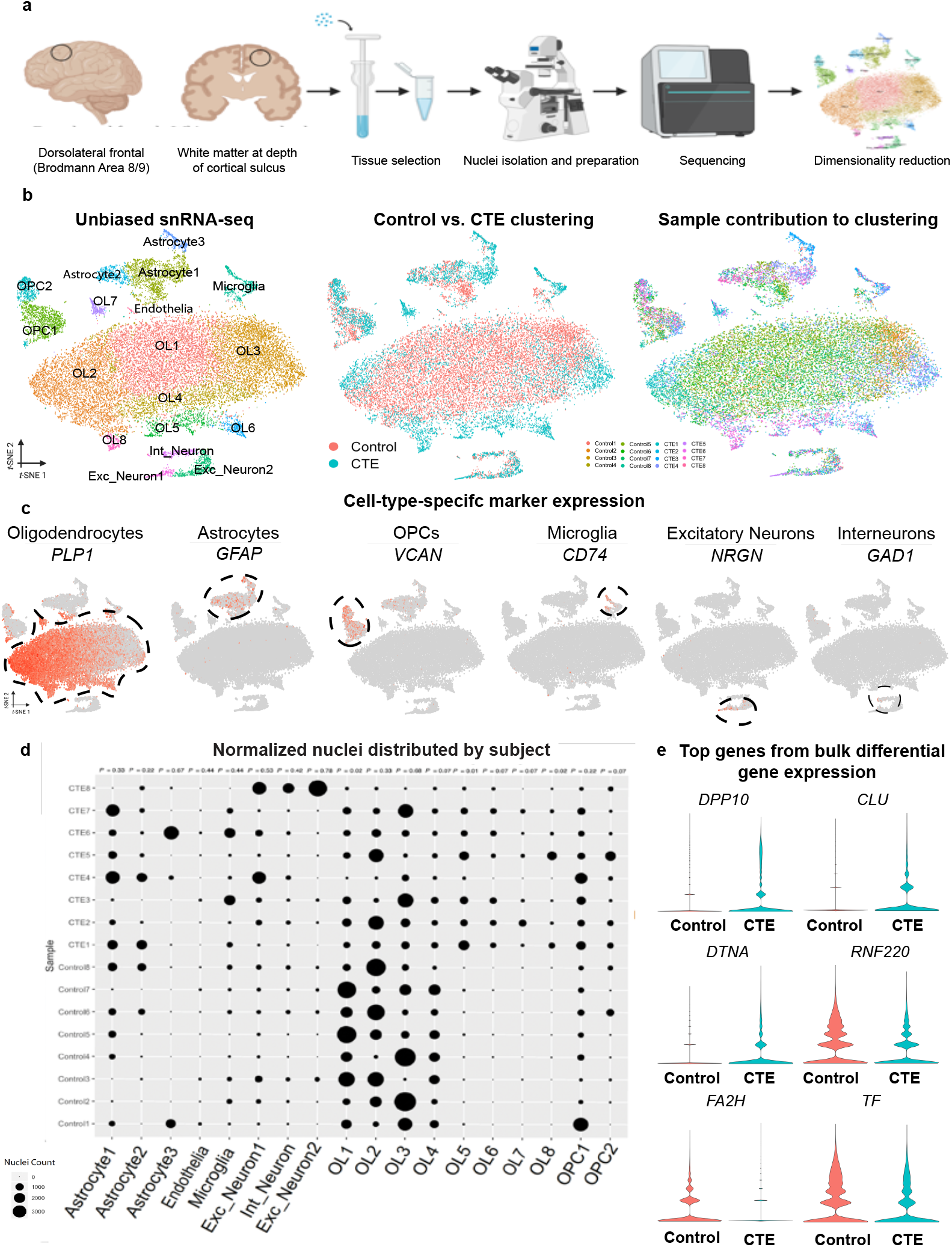
Overview of snRNA-seq and experimental approach. **a.** Dorsolateral frontal cortex area. White matter taken for snRNA-seq analysis and validation. Dounce homogenizer for tissue disassociation. Inverted microscope for single-nucleus hydrogel encapsulation. Sequencer and example data. **b. (left)** Unbiased tSNE showing all major cell-types expected in human brain tissue. **(middle)** tSNE colored by condition, red: Control, teal: CTE. **(right)** tSNE colored by subject, figure key below. **c.** tSNE projection of each major cell type colored by expression of canonical marker genes. Nuclei clusters positive for each marker gene are circled by a dashed line. **d.**Bubble plot of normalized number of nuclei with each subject in the data set on the y-axis and each cell-type and subpopulation on the x-axis. Size of the bubble indicates the number of normalized nuclei in each cell-type for each subject. *P*-values for all nuclei subpopulations listed at top (*n* = 8 per condition; Astrocyte1: *U* = 19.5; Astrocyte2: *U* = 16.5; Astrocyte3: *U* = 27.0; Endothelia: *U* = 22.5; Microglia: *U* = 22.0; Exc_Neuron1: *U* = 24.0; Int_Neuron: *U* = 21.0; Exc_Neuron2: *U* = 29.0; OL1: *U* = 6.0; OL2: *U* = 19.0; OL3: *U* = 27.0; OL4: *U* = 10.0; OL5: *U* = 0.0; OL6: *U* = 5.5; OL7: *U* = 11.0; OL8: *U* = 5.5; OPC1: *P* = 0.22, *U* = 16.0; OPC2: *P* = 0.07, *U* = 11.0; FDR-corrected two-tailed Mann-Whitney U test). **e.** Violin plots of log-normalized counts for top expressed genes identified by bulk differential gene expression between conditions.

We collected intact nuclei by mechanical and enzymatic dissociation and density gradient ultracentrifugation from 1 cm ^3^, frozen white matter tissue blocks harvested immediately beneath the cortical ribbon at the sulcal depth in order to target the subcortical U-fibers, in which pathological abnormalities and axonal microstructural alterations have been reported for CTE^1,8^ (Fig. 1a, Extended Data Fig. 1a). For in situ validations, we collected tissue directly adjacent to the sample taken for snRNA-seq transcriptomic analysis (Extended Data Fig. 1a).

After collecting cell nuclei, we used the inDrops method to encapsulate each nucleus for sequencing. Following sequencing, we performed quality control, cell filtering, and computational analysis using the Seurat pipeline^18,19^ to produce single-nucleus transcriptomic profiles of the 8 control and 8 CTE samples (Fig. 1a,b, Supplementary Table 1, Methods). We report a total of 24,735 droplet-based snRNA-seq profiles across 16 subjects with a median of 561 genes and 854 unique molecular identifiers (UMIs) per nucleus (Fig. 1, Extended Data Fig. 1e-g, Supplementary Table 1). We identified an integrated t-distributed stochastic neighbor embedding (tSNE) projection yielding 18 transcriptomically distinct nuclei clusters derived from control and CTE samples (Fig. 1b, left). We mapped clusters to cell types by identifying the cluster-specific presence of transcripts encoding specific canonical cell-identity markers (Fig. 1c, Extended Data Fig. 2, Supplementary Table 2, Methods). Combining control and CTE subjects, we identified a total of 18,491 oligodendrocytes (~75%), 2,598 astrocytes (~11%), 1,879 OPCs (~8%), 596 microglia (~2%), 857 excitatory neurons (~3%), 271 interneurons (~1%), and 43 endothelial cells (~0.2%). This distribution is similar to what has been previously observed in human white matter tissue^15^.

Examination of the tSNE projection anchored with cell-identity markers suggested multiple differences in CTE as compared to control samples when split by condition (Fig. 1b, middle), including fewer OLs, an increase in the number of disease-specific OL subpopulations, and a shift in the expected astrocyte subpopulation heterogeneity. To determine whether these observed differences between CTE and control tissue partially reflected subject-specific variation rather than condition, we normalized nuclei numbers to the subject with the highest number of nuclei (see methods in^16^) and compared the distributions of total nuclei in each cluster by subject (Fig. 1d). While we did observe subject-specific variation in the proportion of nuclei comprising each cluster, we nonetheless were able to identify statistically significant differences between CTE and control conditions. Specifically, we observed the presence of OL subpopulations that were significantly elevated in CTE, which may suggest a disease-specific OL population^15,20^.

We also conducted bulk gene expression analysis to determine if we could identify a more general disease-specific gene expression signature in CTE as compared to control. Bulk gene expression revealed several genes previously implicated in neurodegeneration. We identified a greater abundance of transcripts encoding Dipeptidyl Peptidase 10 (DPP10), a voltage-gated potassium channel protein positively associated with p-tau-positive neurofibrillary tangles^21^; Clusterin (CLU), a glycoprotein positively associated with amyloid beta (Aβ) toxicity and cell stress^22^; and Dystrobrevin Alpha (DTNA), an astrocytic protein associated with dementia status and p-tau aggregation^23^. We identified lower levels of transcripts encoding RING Finger Protein 220 (RNF220), a protein associated with cholesterol metabolism and neurodegeneration^24^) (Fig. 1e). We also identified lower levels of transcripts encoding Transferrin (TF) and Fatty Acid 2-hydroxylase (FA2H), two proteins associated with iron accumulation^25^ and iron-related neurological diseases^26^ (Fig. 1e).

### Molecular profiling points to fewer OLs in CTE white matter compared to control

To investigate cell-type-specific differences in OLs in CTE compared to controls, we isolated OL nuclei clusters from our primary tSNE projection and re-projected OL nuclei as a subset tSNE by identifying each nucleus previously identified as an OL and performing unbiased graph-based clustering to produce an OL-specific tSNE^14,27^ (Fig. 2a). Consistent with our primary tSNE projection, we observed a statistically significant decrease in the normalized number of OLs in CTE subjects compared to control subjects (*P* = 0.01, two-tailed Mann-Whitney U test) as well as the presence of a CTE-specific OL subpopulation (Fig. 2b, Supplementary Table 5, Methods). These findings suggest OL cell death and myelin loss, consistent with what has been found at the light microscopic level in CTE^1,2,6,8^.

**Figure 2.**
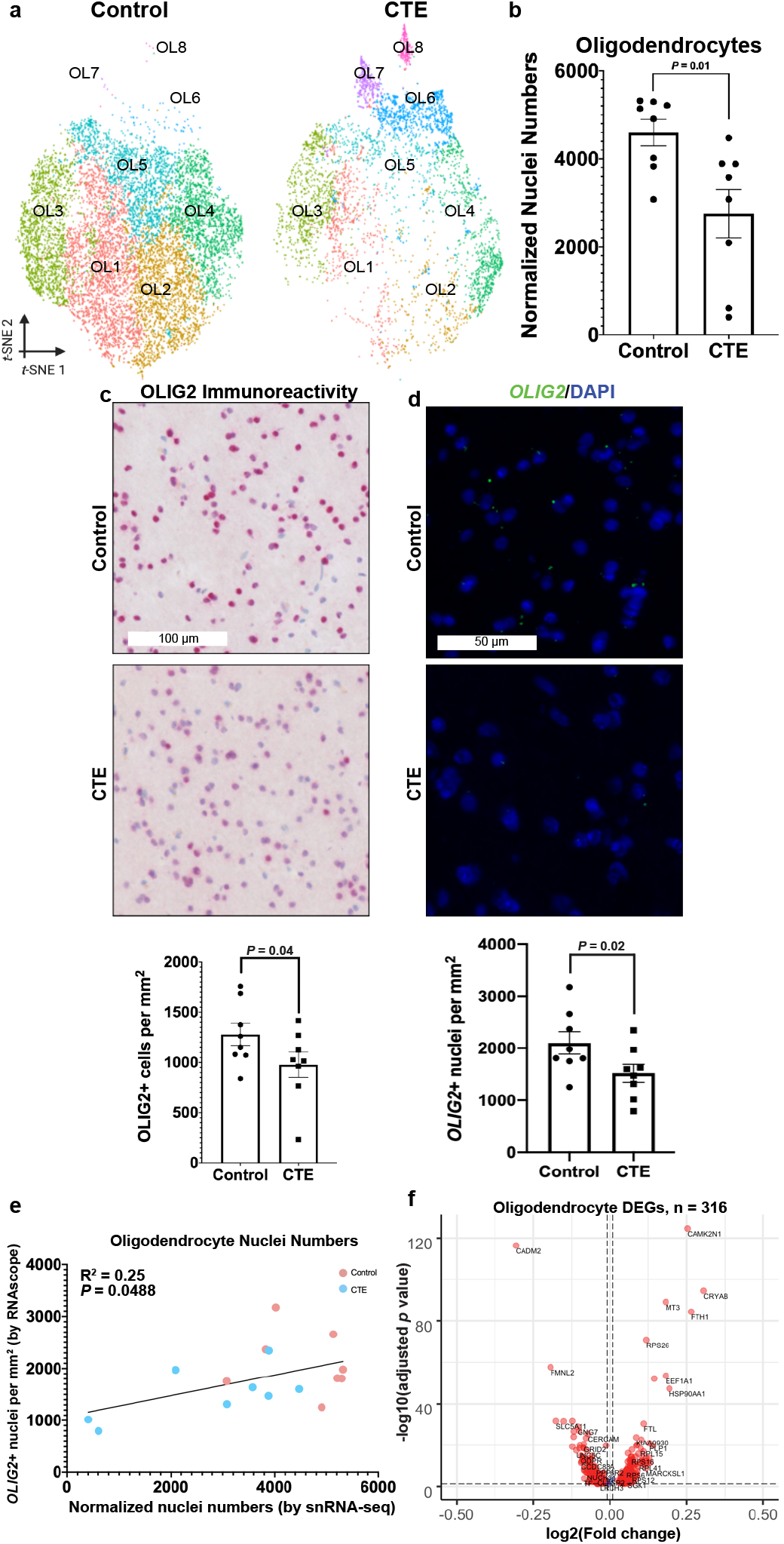
Fewer OLs in CTE white matter compared to control. **a.** tSNE projection of oligodendrocyte nuclei subset from primary tSNE (Fig. 1b). **b.** Scatter plot with bar for normalized nuclei counts for all oligodendrocyte lineage nuclei, dot represents the total normalized number of oligodendrocyte lineage nuclei in one subject (*n* = 8 per condition, median of CTE = 3,331, median of control = 5,023, *P* = 0.01, FDR-corrected two-tailed Mann-Whitney U test, *U* = 9.0). **c.** Representative immunohistochemistry images (anti-OLIG2 for oligodendrocyte lineage cells) of control and CTE tissue. Scale bar **(top, white)**: 100 μm. Quantification **(bottom)** for number of OLIG2-positive nuclei per mm^2^ over the entire white matter sample for each subject (*n* = 8 per condition, *P* = 0.0484, one-tailed Student’s t-test, t(14) = 1.780). **d.** Representative smFISH images for *OLIG2* (green) and DAPI (blue). Scale bar **(top, white)**: 50 μm. Quantification **(bottom)** for number of *OLIG2*-positive nuclei for each subject (*n* = 8 per condition, *P* = 0.03, one-tailed Student’s t-test, t(14) = 2.103). **e.** Scatter plot for number of oligodendrocytes identified by smFISH and snRNA-seq. Each dot represents one subject. Control: red, CTE: teal. (r =.4995, *P* = 0.0488, R^2^ = 0.25). **f.** Volcano plot for differentially expressed oligodendrocyte genes between conditions (Bonferroni correction for multiple comparisons). Vertical dashed lines indicate log2(Fold change) cutoff of 0.01. Horizontal dashed lines indicate −log10(adjusted *P*-value) cutoff of 0.05. Red dots are CTE-associated differentially expressed oligodendrocyte lineage genes (*n* = 316). All data was presented in Mean ± S.E.M.

We performed IHC and smFISH to compare molecular expression in OLs in CTE versus control (Fig. 2c,d, Extended Data Fig. 3). By IHC, we observed fewer OLs (as defined by detection of either Oligodendrocyte Transcription Factor 2 protein (OLIG2)), in CTE compared to control sections, consistent with the transcriptomic findings (*P* = 0.04, one-tailed Student’s t-test) (Fig. 2c, Extended Data Fig. 3a,b). By smFISH, we also observed fewer *OLIG2* positive cells in CTE versus control (*P* = 0.02, one-tailed Student’s t-test) (Fig. 2d, Extended Data Fig. 3c,d). Furthermore, we determined that there were fewer OLs throughout the entire subcortical white matter in CTE, not restricted to the subcortical U-fiber area (Fig. 2c, Extended Data Fig. 3a,b). Notably, we observed a significant, positive correlation between the number of OLs identified by snRNA-seq and smFISH when tracked by subject (Pearson’s r = 0.4995, *P* = 0.0488) (Fig. 2e), suggesting that the normalized nuclei counts accurately reflected in situ cell type abundance.

The in situ probes for OLs also identify OPCs, however, we did not observe differences in OPC numbers in our transcriptomic analysis (*P* = 0.13, two-tailed Mann-Whitney U test) (Extended Data Fig. 4a,b). Furthermore, we performed IHC with anti-PDGFRα, an OPC marker, and found no detectable differences (*P* = 0.23, one-tailed Student’s t-test) (Extended Data Fig. 4c). This suggests that mature OLs are reduced in CTE.

We also analyzed genes differentially expressed in CTE OLs versus controls. This revealed 316 genes with differential transcript levels between conditions, including: Alpha-Crystallin B Chain (*CRYAB* mRNA), an anti-apoptotic and neuroprotective factor^28^, and Ferritin Heavy Chain *(FTH1* mRNA), which is critical for intracellular iron storage^29,30^ (Fig. 2f, Supplementary Table 3).

These analyses suggest that there are fewer OLs in CTE compared to control, supporting OL death and myelin loss. Further, they suggest a potential role for iron accumulation and apoptosis as contributing factors to CTE pathology.

### Iron accumulation and markers of cellular stress response characterize a specific OL subgroup in CTE

Having observed aberrant OL subpopulations in CTE, we examined whether these differences were disease-specific. We first considered the contrast of fewer overall OLs to the apparent increases in OL cell numbers in certain subpopulation clusters in CTE versus controls (Fig. 3a). To determine whether these OL subpopulations consistently harbored more cells in CTE samples, we compared normalized nuclei counts between CTE and control (Fig. 3b, Supplementary Table 5, Methods). All 3 CTE-specific OL groups (OL6, OL7, and OL8) were significantly elevated in CTE compared to control (OL6: *P* = 0.001, OL7: *P* = 0.01, OL8: *P* = 0.05, two-tailed Mann-Whitney U test). We found that these clusters were likely driving the observed greater abundance of transcripts involved in iron accumulation and cellular stress responses (Supplementary Table 2). Differential gene expression analysis between conditions revealed an increase in transcripts encoding iron storage proteins (FTH1 and Ferritin Light Chain (FTL)^29,30^) and CRYAB^28^ (Fig. 3c, Supplementary Table 3), identifying OL6 as the primary subpopulation responsible for the upregulation of these transcripts (Fig. 3d, Supplementary Table 3).

**Figure 3.**
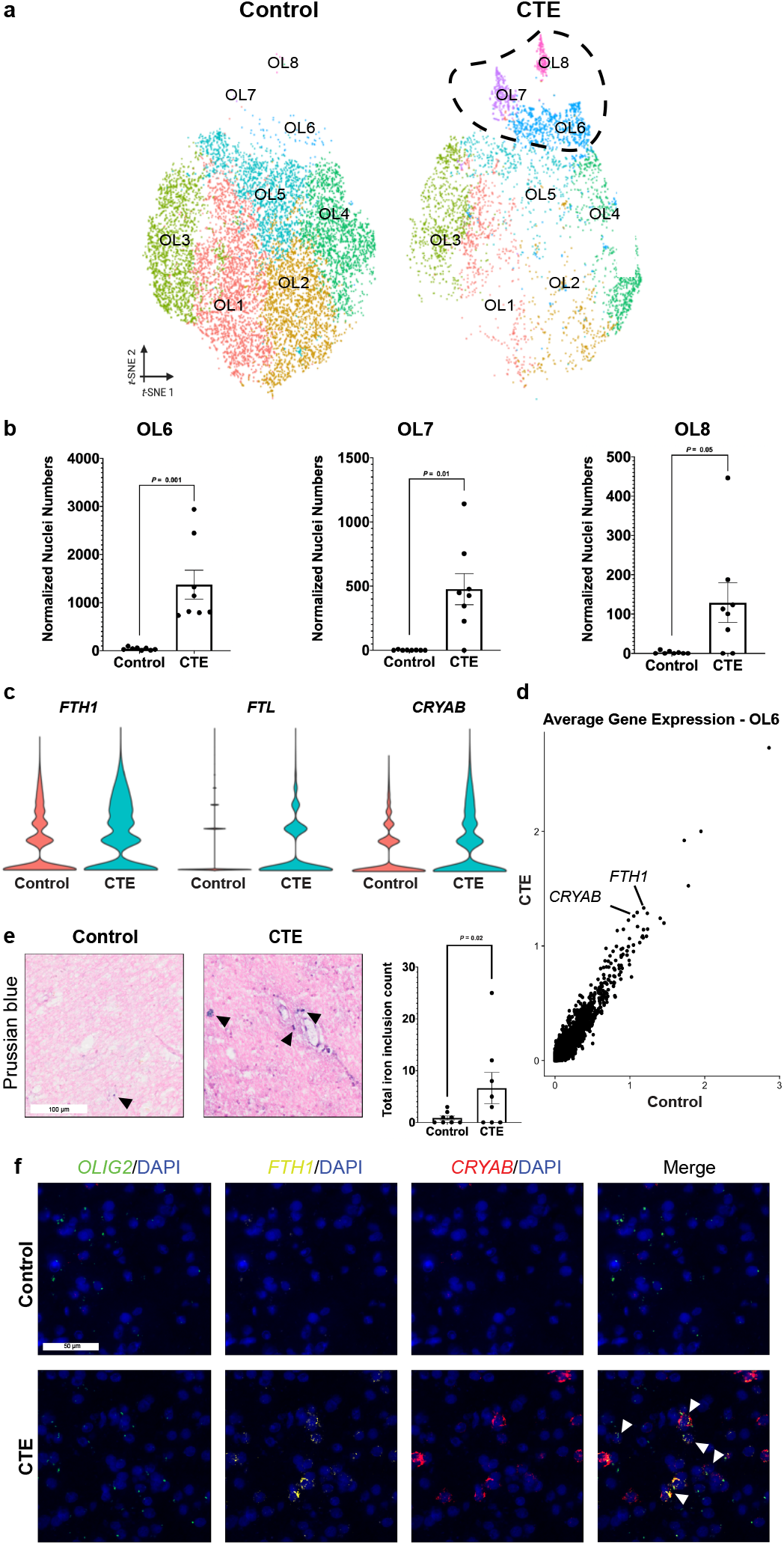
Iron accumulation and markers of cell stress characterized OLs in CTE. **a.** tSNE projection of mature oligodendrocyte lineage nuclei subset from primary tSNE from Fig. 1b with CTE-specific subpopulations circled. **b.** Scatter plot with bar for normalized nuclei counts for CTE-specific oligodendrocyte subpopulations, each dot represents one subject (*n* = 8 per condition; OL6: *P* = 0.001, *U* = 0.0; OL7: *P* = 0.01, *U* = 5.5; OL8: *P* = 0.05, *U* = 11.0; FDR-corrected two-tailed Mann-Whitney U test). **c.** Violin plots of log-normalized counts for top expressed genes in CTE-specific oligodendrocytes. **d.** Scatter plot for average gene expression in OL6. Control gene expression across the x-axis, CTE gene expression across the y-axis. *FTH1* and *CRYAB* are labeled as top differentially expressed genes between conditions. **e.** Representative images of Prussian blue staining for control and CTE tissue. Black arrows label iron inclusions. Scale bar **(left, white)**: 100 μm. Quantification **(right)** for total iron inclusion count, each dot represents a subject (*n* = 8 per condition, *P* = 0.04, one-tailed Student’s t-test, t(14) = 1.878). **f.** Representative multiplexed smFISH images for *OLIG2* (green), *FTH1* (yellow), *CRYAB* (red), and DAPI (blue). *OLIG2* and DAPI **(left)**, *FTH1* and DAPI **(middle-left)**, *CRYAB* and DAPI **(middle-right)**, and merge of all probes **(right)**. White arrows label nuclei positive for *OLIG2*, *FTH1*, *CRYAB*, and DAPI. Scale bar **(top-left, white)**: 50 μm. All data was presented in Mean ± S.E.M.

To investigate iron accumulation in CTE, we analyzed the tissue histochemically for iron aggregates (hemosiderin)^29,31^ (Fig. 3e, Extended Data Fig. 5a). We observed an increase in the total number of iron aggregates in CTE white matter compared to control (*P* = 0.04, one-tailed Student’s t-test). Finally, to validate the presence of OLs with abundant *FTH1* and *CRYAB* transcripts, we performed multiplexed smFISH on control and CTE subjects (Fig. 3f, Extended Data Fig. 5c). We identified sparse groups of *OLIG2* positive nuclei in CTE that were positive for *FTH1* and *CRYAB*, consistent the OL6 subpopulation (Fig. 3f). Together, these findings suggest that aberrant iron accumulation in CTE may result in OL iron accumulation, resultant cellular stress responses, and death.

### Identification of neuroinflammatory astrocytes in CTE white matter

The transcriptomic data also pointed to alterations in the heterogeneity of astrocytes, albeit without changing the overall cell number (Extended Data Fig. 6a). In many neurological diseases, the total number of astrocytes is increased and “reactive” astrocytes predominate^16,32–34^. However, we did not see a difference in the total number of astrocytes in CTE compared to control^2^ (*P* = 0.29, two-tailed Mann-Whitney U test) (Extended Data Fig. 6b). We confirmed this finding by IHC and smFISH (*P* = 0.44 and *P* = 0.50, respectively; one-tailed Mann Whitney U-test, FDR-corrected) (Extended Data Fig. 6c,d).

To further explore CTE-specific astrocyte heterogeneity, we re-projected each astrocyte nucleus data point as a subset tSNE from the primary tSNE as described above for OLs (Fig. 4a) and observed two CTE-specific astrocyte subpopulations, designated Astrocyte2 and Astrocyte3 (Fig. 4a). We found a significant decrease in the number of normalized Astrocyte1 nuclei and a significant increase in the number of normalized Astrocyte2 and Astrocyte3 nuclei in CTE as compared to controls (Astrocyte1: *P* = 0.02, Astrocyte2: *P* = 0.01, Astrocyte3: *P* = 0.03, two-tailed Mann-Whitney U test, FDR-corrected) (Fig. 4b, Supplementary Table 5, Methods). To reveal the transcripts driving the emergence of each subpopulation, we performed cell-type marker analysis (Supplementary Table 2). The Astrocyte1 group expressed many genes associated with normal functioning astrocytes (e.g. *GFAP,* Catenin Delta-2 (*CTNND2*), Aquaporin-4 (*AQP4*)), while the Astrocyte2 group expressed transcripts associated with dysfunctional metabolism (e.g. Pyruvate Dehydrogenase Kinase 4 (*PDK4*)^35^), and the Astrocyte3 group showed enrichment for transcripts associated with neuroinflammation and aging (Cluster of Differentiation 44 (*CD44*), B-Cell Lymphoma 6 (*BCL6*), and Alpha 1-antichymoptrypsin (*SERPINA3*)^16,33^) (Fig. 4c, Supplementary Table 3). Additional gene expression analysis for the Astrocyte3 group transcript profile confirmed increases in mRNAs related to neuroinflammation, suggesting that Astrocyte3 may represent a CTE-specific astrocyte subpopulation and/or neuroinflammatory state (Fig. 4d, Supplementary Table 3).

**Figure 4.**
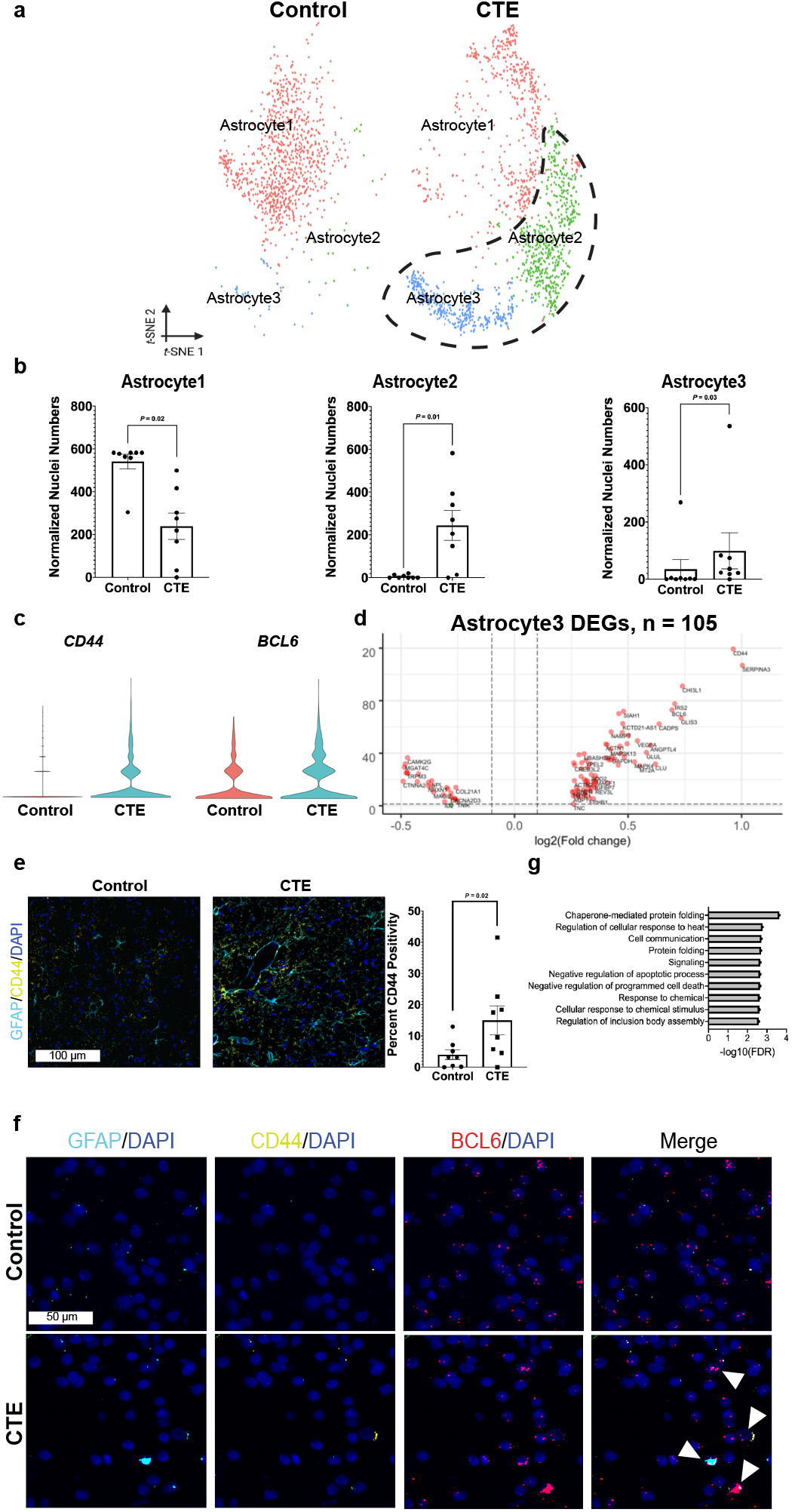
Neuroinflammatory astrocytes in CTE white matter. **a.** tSNE projection of astrocyte nuclei subset from primary tSNE (Fig. 1b). **b.** Scatter plot with bar for normalized nuclei counts for astrocyte subpopulations, each dot represents one subject (*n* = 8 per condition; Astrocyte1: *P* = 0.02, *U* = 2.0; Astrocyte2: *P* = 0.01, *U* = 7.5; Astrocyte3: *P* = 0.03, *U* = 12.0; FDR-corrected two-tailed Mann-Whitney U test). **c.** Violin plots of log-normalized counts for top expressed genes in CTE-specific astrocytes. **d.** Volcano plot for differentially expressed Astrocyte3 genes between conditions (Bonferroni correction for multiple comparisons). Vertical dashed lines indicate log2(Fold change) cutoff of 0.01. Horizontal dashed lines indicate −log10(adjusted *P*-value) cutoff of 0.05. Red dots are CTE-associated differentially expressed Astrocyte3 genes that pass both cutoffs. *n* = 105 differentially expressed genes. **e.** Representative IF images for GFAP (teal), CD44 (yellow), and DAPI (blue) in control and CTE. Scale bar **(left, white)**: 100 μm. Quantification **(right)** for percent of total area positive for CD44 IF for each subject (*P* = 0.02, one-tailed Student’s t-test). **f.** Representative multiplexed smFISH images for *GFAP* (teal), *CD44* (yellow), *BCL6* (red), and DAPI (blue). *GFAP* and DAPI **(left)**, *CD44* and DAPI **(middle-left)**, *BCL6* and DAPI **(middle-right)**, and merge of all probes **(right)**. White arrows label nuclei positive for *GFAP*, *CD44*, *BCL6*, and DAPI. Scale bar **(top-left, white)**: 50 μm. **g.** Top 10 enriched gene ontology terms ranked by FDR. All data was presented in Mean ± S.E.M.

To validate finding more astrocytes with enriched levels of *CD44* transcripts in CTE compared to control, we assessed in situ CD44 protein by IF, co-staining with the astrocyte marker GFAP (Fig. 4e, Extended Data Fig. 7a). We observed a significant increase in total area of CD44 immunopositivity in CTE compared to control (*P* = 0.02, one-tailed Student’s t-test) (Fig. 4e, right), consistent with an increase in the number of astrocytes expressing *CD44* and an increase in the Astrocyte3 cluster. Further, by multiplexed smFISH, we found transcript profiles consistent with the Astrocyte3 cluster in CTE (*CD44* and *BCL6*) (Fig. 4f, Extended Data Fig. 7b).

We also performed gene ontology (GO) analysis^36^ using the identified differentially expressed genes in astrocytes in CTE versus controls. We found a significant increase in transcripts related to processes such as protein folding, heat response, and regulation of apoptosis, which are often observed in disease-associated astrocytes^33^ (Fig. 4g, Supplementary Table 4).

Together, these findings suggest that the overall number of astrocytes in CTE white matter is unchanged compared to controls, but there is a shift to greater astrocyte subpopulations demonstrating neuroinflammatory and dysfunctional metabolic states. These findings support the possibility that astrocytes in CTE alter their genetic profiles to become neuroinflammatory.

## DISCUSSION

We performed a transcriptomic analysis of dorsolateral frontal white matter from 16 male subjects, 8 with neuropathologically-verified CTE and 8 age- and PMI-matched controls and report the analyses of 24,735 cell nuclei. We found fewer oligodendroglia in CTE compared to controls and differences in oligodendroglial and astrocyte subpopulations. The transcriptomically unique profiles distinguishing OL subtypes and/or OL cell states in CTE point to an enrichment of phenotypes involving cell stress response and iron accumulation (*CRYAB*^28^ and *FTH1* transcripts, respectively^29,30^). With respect to astrocytes, transcriptomic analyses suggest enrichment of cell subtypes and/or states associated with neuroinflammation and dysfunctional metabolism, without a change in overall astrocyte number. We did not observe transcriptional or cell-type differences associated with neurons or microglia, likely due to under sampling of those cell nuclei.

Neuroinflammation, blood brain barrier leakage, and hemosiderin-laden macrophages are characteristic of the white matter in CTE^3,6,37,38^. Iron accumulation induces transcription of the ferritin genes *FTH1* and *FTL*^39^ and also activates ferroptosis, a form of cell death characterized by iron-dependent accumulation of lipid hydroperoxides^40^. The data reported here and validated across multiple dimensions– snRNA-seq, smFISH, and IHC/IF–suggest a role for OLs in the pathogenesis of CTE. Our data also suggest that OL loss in CTE might occur through accumulation of excess iron, as reflected by increased iron histologically, enrichment of transcripts encoding the intracellular ferritin proteins FTH1 and FTL, and OL cell death, evidenced by reduced cell number and more abundant levels of transcripts encoding the apoptosis and protein aggregation regulator, CRYAB. In addition, the observed lower levels of transcripts encoding TF (which sequesters iron from brain blood vessels^41^) and FA2H (which is implicated iron-related neurological disorders and axonal and myelin sheath integrity^26,42–44^) in CTE support widespread iron trafficking and aggregation abnormalities.

The presented datasets serve as a mineable resource for understanding the molecular and cellular dysregulation associated with CTE. Integration with additional data sets will enable expansion of CTE transcriptomic profiles to further characterize potential mechanisms of CTE pathogenesis. Additional studies are warranted to determine whether these same changes occur in other brain regions, parallel the progression of p-tau pathology, or correlate with symptom severity in CTE. In any case, the present findings provide a catalyst for the future development of biomarkers to aid in the diagnosis of CTE during life as well as potential targets for therapeutics.

## METHODS

### Human patient selection

CTE subjects were selected based on a neuropathologically verified diagnosis of CTE Stage II or Stage III by A.C.M., T.D.S., B.R.H. or V.E.A. without any co-morbid neurodegenerative disease and availability of fresh frozen from dorsolateral frontal cortex (Brodmann area 8/9) from the Veterans Affairs-Boston University-Concussion Legacy Foundation (VA-BU-CLF)/ Understanding Neurological Injury and Traumatic Encephalopathy (UNITE) Brain Bank. Control subjects were age-, PMI-, RIN-, and sex-matched (all males) with neuropathological confirmation of the absence of neurological or neurodegenerative disease and availability of fresh frozen tissue from dorsolateral frontal cortex (BA 8/9) from the National Posttraumatic Stress Disorder (PTSD) Brain Bank. For the snRNA-seq analysis, the subcortical white matter was dissected from the tissue blocks and included both U-fibers and deep white matter. For the in situ analysis, a section of BA 8/9 tissue with both cortical and white matter tissue adjacent to the white matter block taken for snRNA-seq analysis was isolated. Subject ages were not significantly different (Control: 55.6 ± 7.5 years; CTE: 55.9 ± 10.7 years; *P* = 0.94, two-tailed Mann-Whitney U test). Subject PMI was not significantly different (Control: 31.7 ± 9.7 hours; CTE: 35.3 ± 10.2 hours; *P* = 0.70, two-tailed Mann-Whitney U test). Subject RIN values were not significantly different (Control: 6.4 ± 1.7; CTE: 6.8 ± 1.7; *P* = 0.56, two-tailed Mann Whitney U test). RIN number determination was completed according to manufacturer instructions using a Bioanalyzer RNA 6000 Nano kit (Agilent).

### Isolation of nuclei from frozen post-mortem human brain tissue

All tissue was macrodissected into 30 mg sections from dorsolateral frontal white matter (BA 8/9) with a scalpel blade and stored at −80° Celsius. On the day of single-nucleus encapsulation, tissue was placed in a 7 mL dounce with ice cold buffer (0.5M sucrose, 2M KCl, 1M MgCl2, 1M Tricine-KOH pH 7.8, spermine, spermidine, DTT, RNasin, H2O) and dounced with a tight pestle ten times to mechanically dissociate and homogenize the tissue. To further dissociate the tissue, 5% IGEPAL-CA630 was added to the homogenate and dounced 5 times. The homogenate was then strained through a 40 μm cell strainer (VWR) on ice. 5 mL of 50% iodixanol was added to the homogenate and mixed. In a 13 mL ultraclear ultracentrifuge tube, 1 mL of 40% iodixanol was added with 1mL of 30% iodixanol carefully layered on top, followed by 10 mL of the tissue homogenate. The tissue was then ultracentrifuged in a swing-bucket rotor at 10,000xg for 18 minutes. Following ultracentrifugation, the 35% iodixanol layer was collected and the nuclei were counted. Nuclei were diluted with 30% iodixanol to 90,000 nuclei/mL for inDrops chip loading. Nuclei were isolated for inDrops using a previously described method^45^.

### Droplet-based snRNA-seq with inDrops and sequencing

Isolated nuclei were delivered to the inDrops single cell core facility at Harvard Medical School. The inDrops core facility generated inDrops V3 nuclei libraries for each subject by encapsulating each nucleus in a hydrogel containing barcodes and unique molecular identifiers (UMIs). Nuclei were lysed and each free mRNA molecule was reverse transcribed to cDNA and labeled with a barcode and UMI, which enables mapping that specific cDNA molecule to its particular cell of origin and ensures that specific amplified sequence is counted only once bioinformatically^46,47^, thus avoiding the production of potential PCR amplification-generated artifact. This process produced labeled cDNA molecules for all 16 samples. Sample cDNA was delivered to the Biopolymers Facility at Harvard Medical School. Sample quality was assessed by Agilent Tapestation. Samples with excess primer dimer contamination were filtered by SPRIselect size selection (Beckman Coulter, B23317), followed by pooling of libraries and dilution for sequencing. Pooled samples were sequenced on the Illumina NextSeq500 at an average of 40,000 reads per nucleus barcode, approximately 360,000,000 reads per sample.

### Pre-processing snRNA-seq sequencing data

The Biopolymers Core Facility delivered Binary Base Call (BCL) files for each sequencing run. The inDrops single cell method requires four primers to completely capture and de-multiplex a cDNA molecule. The first primer captures the transcript, the second captures part of the single cell barcode, the third captures the library or sample index, and the fourth captures the rest of the single cell barcode, the UMI, and the polyT tail. BCL files produced by sequencing were converted to 4 FastQ files, one per inDrops primer read. FastQ files were de-multiplexed by first filtering reads, only keeping reads that followed the expected inDrops index structure, described above. Reads that passed structure filtering were de-multiplexed based on library index, resulting in reads by sample of origin, i.e., subject. Abundant barcodes were identified and sorted so that each cDNA from a single nucleus was grouped together. After each read was de-multiplexed, thus assigning reads to nucleus of origin and nuclei to sample of origin, reads were aligned with bowtie/1.1.1 to an edited version of the GRCh38.97 reference genome so that all exonic and intronic reads were counted. Aligned reads were quantified and a UMI count matrix was produced for each sample. De-multiplexing was executed following the inDrops analysis pipeline (https://github.com/indrops/indrops).

### Quality control and cell filtering

UMI count matrices for each subject were loaded into R version 3.6.1 using the Scater R package^48^. Control samples were combined and CTE samples were combined to generate two separate objects with the R package Seurat version 3.1^18^. The Seurat object filtration process included excluding nuclei with fewer than 400 genes and more than 2,500 genes, which limited our analysis to nuclei with high-quality RNA. We also removed nuclei with more than 1% mitochondrial genes, which excluded nuclei that were potentially undergoing lysis due to sample preparation. To identify cell types and/or cell states in the data set, we constructed an unbiased *k*-nearest neighbor graph based on top principal components followed by a Jaccard similarity step and the Louvain modularity optimization algorithm^49^ to group the cell nuclei into clusters based on gene expression similarities and produce a two-dimensional t-distributed stochastic neighbor embedding (tSNE) projection^18,50^. The tSNE algorithm provides an unbiased similarity clustering of nuclei by gene expression, shaped by parameters selected prior to analysis, such as number of principal components, number of highly variable genes, and resolution^27^. Through iterative parameter space testing^13–17,27^, we settled on values that resulted in stable clustering with high intercluster distance and low intracluster distance, meaning, on average, nuclei in the same cluster were more similar to each other than to nuclei in a different cluster. Specifically, we chose 23 principal components and a resolution of 0.8 for input parameters (Extended Data Fig. 1h). The Seurat 2 objects were then normalized and log-transformed using Seurat’s implementation of SCTransform and integrated using Seurat’s implementation of canonical correlation analysis^18,19^ (CCA). Integration was performed to integrate control and CTE samples, thus creating a single Seurat object for downstream analyses, “CTE.combined.”

### Cell clustering

The single integrated Seurat object, “CTE.combined,” was scaled using ScaleData and a principal component analysis (PCA) was performed for initial dimensionality reduction. An Elbow Plot was produced to determine the number of principal components to be used for further analyses (Extended Data Fig. 1h). Based on the Elbow Plot, 23 principal components were used. Using RunTSNE, FindNeighbors, and FindClusters, “CTE.combined” was further reduced in dimensionality to produce a two-dimensional t-distributed stochastic neighbor embedding (tSNE) projection. This process included constructing an unbiased *k*-nearest neighbor graph based on top principal components followed by a Jaccard similarity step and the Louvain modularity optimization algorithm. Parameters, including resolution, were varied over several iterations to identify a tSNE that was in agreement with published human white matter brain data^15^ and that represented nuclei clusters that were consistently present over several iterations. Nuclei distances were also considered in order to increase intercluster distances and decrease intracluster distances, thereby producing clusters in which nuclei in the same cluster were more similar to each other than to nuclei in a different cluster. The final tSNE projection used was produced at resolution 0.8 and included 18 transcriptionally distinct nuclei clusters.

### Cell-type annotation and sub-clustering

Nuclei clusters were annotated using canonical cell-type marker genes: *PLP1* for oligodendrocytes, *GFAP* for astrocytes, *VCAN* for OPCs, *CD74* for microglia, *NRGN* for excitatory neurons, *GAD1* for interneurons, and *CLDN5* for endothelial cells. FindAllMarkers and FindMarkers were used to identify cluster-specific genes^13–16^. Top genes for each cluster and the expression of top genes on the tSNE were used to identify clusters (Fig. 1c, Extended Data Fig. 2, Supplementary Table 2). To further analyze specific cell-types, clusters from the primary tSNE were subset. SubsetData was used to isolate nuclei from a specific cell-type, and ScaleData, RunPCA, RunTSNE, FindNeighbors, and FindClusters were performed on the subset data to produce a new Seurat object for each cell-type.

### Cell-type-specific gene expression analysis and differential gene expression analysis

To determine the marker genes (the genes that make a specific cluster unique compared to the other clusters) in any given cluster or subset cluster, the Seurat function FindMarkers was used. Differential gene expression analysis was performed using Seurat’s implementation of MAST^51^. For bulk analysis, all CTE clusters were analyzed against all control clusters. For cell-type-specific analysis, one cluster from CTE was analyzed against the corresponding cluster from control. Bonferroni-corrected *P*-value < 0.05 was taken, and genes with a fold change of at least 0.01 were considered.

### Technique correlation analysis

To determine whether findings across experimental methods were significantly correlated, normalized nuclei counts identified by snRNA-seq transcriptomic analysis were plotted against the number of *OLIG2*+ nuclei identified by smFISH. Each dot on the plot represented a single human subject (Control: Red, CTE: Teal), with snRNA-seq transcriptomic analysis providing the x-coordinate value and smFISH providing the y-coordinate value. A Shapiro-Wilk test was performed to determine normality of the data set, and a Pearson correlation coefficient was produced to determine correlation and statistical significance.

### Gene ontology analysis

The set of all genes significantly differentially expressed between CTE and control nuclei (FDR < 0.05) within astrocyte cell-types were used as query inputs for gene ontology analyses by PANTHER Classification System v14.1 (www.pantherdb.org)^36^ to identify over-represented gene modules. Modules were assigned by PANTHER's statistical overrepresentation test in which a binomial test was applied to each query set relative to its corresponding reference gene set (all genes expressed in a certain cell-type) to determine if a known biological pathway had non-random linkage within the query set. All identified modules derived from PANTHER’s “GO biological process complete” gene annotation and only those with a false-discovery-rate-corrected *P*-value < 0.05 were considered.

### Cell-type representation analysis

To determine whether any cluster of nuclei was increased or decreased in CTE subjects, compared to control subjects, numbers of nuclei in each cluster were normalized to the sample with the largest number of nuclei (see methods in ^16^, Supplementary Table 5). The following formulas for normalization were used:

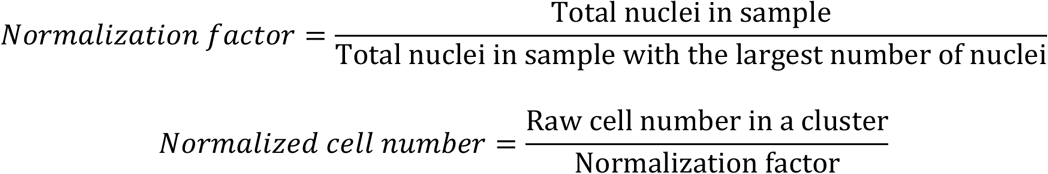

Normalized cell numbers for each cell-type and sample were then compared between conditions using a two-tailed Mann-Whitney U test (FDR-corrected for multiple comparisons).

### Iron staining

Fresh frozen tissue stored at −80° Celsius was adhered to cryostat compatible metal chucks using Optimal Cutting Temperature compound (Leica) and allowed to acclimate to a −17° Celsius cryostat (Leica CM3050 S) for 1 hour before cutting. Tissue was cut at 16 μm thick sections and placed on Fisher Superfrost Plus (Fisher Scientific) slides. All slides were heated in a 60° Celsius oven for 15 minutes after cutting for tissue adherence and placed directly in room temperature 10% formalin for 1.5 hours for fixation. Slides were then dehydrated in 50%, 70%, and 100% ethanol baths for 2 minutes each before being heated in a 60° Celsius oven for 30 minutes to dry slides for room temperature storage. Prussian blue staining, which produces a dark blue pigment on all ferrous iron present in the tissue, was completed on all slides with hematoxylin and eosin counterstain. Prussian blue stained slides were placed in a solution of 0.06N Hydrochloric acid and Potassium Ferricyanide for 1 hour at room temperature before a wash in 1% Acetic Acid and hematoxylin and eosin counterstains.

### Immunofluorescence

Fresh frozen tissue stored at −80° Celsius was adhered to cryostat compatible metal chucks using Optimal Cutting Temperature compound (Leica) and allowed to acclimate to a −17° Celsius cryostat (Leica CM3050 S) for 1 hour before cutting. Slides used for IF staining were cut at 10 μm using a CryoJane Tape Transfer System (Leica). Slides were placed in ice cold formalin for 15 minutes after cutting, rinsed in PBS, and allowed to dry fully under a ventilation hood. Slides were then stored at −80° Celsius and allowed to thaw at 4° Celsius before being used for manual IF staining. Anti-mouse GFAP (1:1,000; Millipore) and anti-rabbit CD44 (1:2,000; Abcam) antibodies were used for double-labeled IF staining. Slides were placed in 3 changes of 0.01% Triton for 5 minutes for tissue permeabilization, blocked in 3% normal donkey serum for 30 minutes, heated at 95° Celsius in AR6 solution (PerkinElmer) for 20 minutes, and placed in a 4° Celsius refrigerator overnight with a primary antibody. A donkey anti-mouse or a donkey anti-rabbit secondary was used at a 1:500 concentration with 1% Normal Donkey Serum for one hour at room temperature. Fluorescent labeling was conducted using Opal 480 (GFAP) and Opal 570 (CD44) dyes (1:150; Akoya Biosciences) for 10 minutes at room temperature, followed by 5 minutes of DAPI counterstaining.

### Immunohistochemistry

Fresh frozen tissue stored at −80° Celsius was adhered to cryostat compatible metal chucks using Optimal Cutting Temperature compound (Leica) and allowed to acclimate to a −17° Celsius cryostat (Leica CM3050 S) for 1 hour before cutting. Tissue was cut at 16 μm thick sections and placed on Fisher Superfrost Plus (Fisher Scientific) slides. All slides were heated in a 60° Celsius oven for 15 minutes after cutting for tissue adherence and placed directly in room temperature 10% formalin for 1.5 hours for fixation. Slides were then dehydrated in 50%, 70%, and 100% ethanol baths for 2 minutes each before being heated in a 60° Celsius oven for 30 minutes to dry slides for room temperature storage. All brightfield immunohistochemistry was completed using individually optimized protocols using Research Detection Kits on a BOND RX (Leica) system. Antibodies used on the BOND RX included: anti-rabbit PDGFRα (1:100; Abcam) and anti-rabbit OLIG2 (1:100; Millipore).

### Single-molecule fluorescent mRNA in situ hybridization

Fresh frozen tissue stored at −80° Celsius was adhered to cryostat compatible metal chucks using Optimal Cutting Temperature compound (Leica) and allowed to acclimate to a −17° Celsius cryostat (Leica CM3050 S) for 1 hour before cutting. Tissue was cut at 16 μm thick sections and placed on Fisher Superfrost Plus (Fisher Scientific) slides. All slides were heated in a 60° Celsius oven for 15 minutes after cutting for tissue adherence and placed directly in room temperature 10% formalin for 1.5 hours for fixation. Slides were then dehydrated in 50%, 70%, and 100% ethanol baths for 2 minutes each before being heated in a 60° Celsius oven for 30 minutes to dry slides for room temperature storage. Single-molecule fluorescent mRNA in situ hybridization was conducted using RNAscope (Advanced Cell Diagnostics) probes and kits optimized for use on a BOND RX (Leica) system. All slides were treated with a 30 minute protease step, 1 minute of hybridization, and 10 minutes of epitope retrieval at the beginning and end of the protocol. Concentration of Opal dyes (480, 570, 690; Akoya Biosciences) for fluorescent labeling were diluted at 1:1,500 in TSA Buffer (Advanced Cell Diagnostics) for every probe, all other steps were completed according to manufacturer's instructions.

### Slide visualization and quantification

All slides used for iron staining, IF, IHC, and smFISH were scanned at 40X using custom protocols on a Vectra Polaris (Akoya Biosciences) whole slide scanner and quantified using custom protocols on HALO (Indica Labs) software. For IHC analysis, HALO was manually trained to identify the color of the antibody-positive chromogen or dye (DAB, fast red, or Prussian Blue) and counterstain (hematoxylin). By manually adjusting HALO settings on stain color identification and expected inclusion shape and size, chromogen- or dye-positive inclusion counts and/or positively staining area measurements were generated for the selected tissue analysis area. Masks representing the HALO analysis were produced in order to determine the accuracy of all analyses. For IF and smFISH, HALO is programmed innately to detect the Opal dyes and DAPI used in these experiments. For IF analysis, HALO settings on Opal dye intensity were adjusted to accurately map observed GFAP and CD44 expression. HALO produced values representing the area of GFAP or CD44 positivity based on the manually adjusted positivity settings in the selected tissue analysis area. For RNAscope analysis, HALO settings on dye intensity and inclusion shape were adjusted to identify and map DAPI positive nuclei. HALO settings on Opal dye intensity were adjusted to accurately encompass Opal dye-positive inclusions inside of nuclei and/or within a 5 μm radius from the center of nuclei. This value was selected to best represent measured cell bodies observed in our OLIG2 IHC and GFAP IF staining. Inclusions detected within the set parameters were measured by area. Masks representing the HALO analysis were produced in order to determine the accuracy of all analyses. All analyses were conducted in 1 mm^2^ sections of the white matter tissue adjacent to the depth of a cortical sulcus unless otherwise specified.

### Statistical analyses

Statistical analyses were performed in either R version 3.6.1 or GraphPad Prism 8. Gene expression statistical analyses were performed in R using MAST and Bonferroni correction for multiple comparisons^51^. To identify an increase or decrease in the number of normalized nuclei between conditions, a Mann-Whitney U test was performed and *P*-values were FDR-corrected for multiple comparisons. IHC, IF, and smFISH quantification was performed using a one-tailed Student’s t-test or one-tailed Mann-Whitney U test. No statistical methods were used to predetermine sample size. Sample size was determined by tissue availability.

## Code availability

Source code used for all computational analyses can be found on GitHub at: https://github.com/blakechancellor/Altered-oligodendroglia-and-astroglia-in-chronic-traumatic-encephalopathy.

## Data availability

The snRNA-seq and in situ validation data that support the findings of this study are available from the corresponding author. Raw and processed snRNA-seq data are available at Gene Expression Omnibus (GEO) under accession number GSE155114.

## ACKNOWLEDGMENTS

We thank the VA-BU-CLF/UNITE Brain Bank and the National PTSD Brain Bank for providing human brain tissue; the Harvard Medical School Single Cell Core and the inDrops core facility for single-nucleus RNA-seq sample preparation; and the Harvard Biopolymers Facility for sample quality control and RNA sequencing. This work was supported in part by the Harvard/MIT Joint Research Grants Program in Basic Neuroscience (K.B.C., J.E.D.-C., A.C.M.) and the Concussion Legacy Foundation Young Investigator Fellowship (S.E.C.). We would like to acknowledge the many families who contributed to this work and the clinical and neuropathology research staff of the BU CTE Center, VA Boston Healthcare System and Edith Nourse Rogers VA Medical Center. This work was supported by grant funding from: NIA (AG057902, AG06234), NINDS (U54NS115266, U01NS086659), National Institute of Aging Boston University AD Center (P30AG13846); the Concussion Legacy Foundation and the Nick and Lynn Buoniconti Foundation. The views, opinions and/or findings contained in this article are those of the authors and should not be construed as an official Veterans Affairs position, policy or decision, unless so designated by other official documentation. Funders did not have a role in the design and conduct of the study; collection, management, analysis, and interpretation of the data; preparation, review, or approval of the manuscript; or decision to submit the manuscript for publication.

## AUTHOR CONTRIBUTIONS

K.B.C., S.E.C., and A.C.M. designed all experiments. B.R.H., T.D.S., V.E.A., and A.C.M. performed neuropathological evaluation of each subject. S.E.C. collected all tissue. K.B.C. performed snRNA-seq sample preparation. With advice from B.W.O. and assistance from J.E.D.-C., K.B.C. performed computational analysis of the snRNA-seq data. S.E.C., with assistance from K.B.C., performed all in situ validation. K.B.C., S.E.C., S.M.D., and A.C.M. wrote the manuscript with input from the co-authors. K.B.C., S.E.C., and A.C.M. oversaw all aspects of the study.

## ADDITIONAL INFORMATION

Correspondence should be addressed to B.W.O., S.M.D., and A.C.M. (

---

**Extended Data Table 1.** Demographic and clinical profiles of subjects.

| Subject | Tissue Origin | Brain Region | Sex | Age | Sport Exposure | RIN | COD | CTE Stage | PMI (hours) |
| --- | --- | --- | --- | --- | --- | --- | --- | --- | --- |
| CTE1 | VA-BU-CLF/UNITE Brain Bank | Brodmann Area 8/9 | Male | 65 | NFL | 6 | Myocardial Infarct | CTE Stage III | 37 |
| CTE2 | VA-BU-CLF/UNITE Brain Bank | Brodmann Area 8/9 | Male | 69 | NFL | 6.9 | Cancer | CTE Stage III | 24 |
| CTE3 | VA-BU-CLF/UNITE Brain Bank | Brodmann Area 8/9 | Male | 52 | NFL | 6.5 | Hepatic Encephalopathy | CTE Stage III | 48 |
| CTE4 | VA-BU-CLF/UNITE Brain Bank | Brodmann Area 8/9 | Male | 70 | NFL, AFL, College Football | 7.3 | Cardiac Event | CTE Stage II | 31 |
| CTE5 | VA-BU-CLF/UNITE Brain Bank | Brodmann Area 8/9 | Male | 38 | NFL | 8.1 | Hypertensive Cardiomyopathy | CTE Stage II | 24 |
| CTE6 | VA-BU-CLF/UNITE Brain Bank | Brodmann Area 8/9 | Male | 49 | College Football, Army, MMA | 2.8 | Gastric Cancer | CTE Stage II | 24 |
| CTE7 | VA-BU-CLF/UNITE Brain Bank | Brodmann Area 8/9 | Male | 47 | College Football | 8.6 | Alcohol-related necrotizing pancreatitis | CTE Stage III | 47 |
| CTE8 | VA-BU-CLF/UNITE Brain Bank | Brodmann Area 8/9 | Male | 57 | USFL | 8 | Cirrhosis | CTE Stage III | 47 |
| Control1 | Posttraumatic Stress Disorder Brain Bank | Brodmann Area 8/9 | Male | 65 | None | 4.4 | Atherosclerotic Cardiovascular Disease | N/A | 27 |
| Control2 | Posttraumatic Stress Disorder Brain Bank | Brodmann Area 8/9 | Male | 60 | None | 7.7 | Saddle Pulmonary Thromboembolism & Deep Venous Thrombosis | N/A | 30 |
| Control3 | Posttraumatic Stress Disorder Brain Bank | Brodmann Area 8/9 | Male | 53 | None | 5.1 | Multiple Injuries | N/A | 35.5 |
| Control4 | Posttraumatic Stress Disorder Brain Bank | Brodmann Area 8/9 | Male | 69 | None | 5.7 | Suicide | N/A | 38 |
| Control5 | Posttraumatic Stress Disorder Brain Bank | Brodmann Area 8/9 | Male | 48 | None | 9.5 | Accidental Traumatic Asphyxiation | N/A | 10.5 |
| Control6 | Posttraumatic Stress Disorder Brain Bank | Brodmann Area 8/9 | Male | 51 | None | 7.7 | Hypertensive Atherosclerotic Cardiovascular Disease | N/A | 29 |
| Control7 | Posttraumatic Stress Disorder Brain Bank | Brodmann Area 8/9 | Male | 48 | None | 7 | Morbid Obesity Complications | N/A | 45 |
| Control8 | Posttraumatic Stress Disorder Brain Bank | Brodmann Area 8/9 | Male | 51 | None | 4.4 | Asphyxia | N/A | 38.5 |

Extended Data Table 1. Demographic and clinical profiles of subjects.
| Abbreviations |  |
| --- | --- |
| NFL | National Football League |
| AFL | American Football League |
| MMA | Mixed Martial Arts |
| USFL | United States Football League |
| RIN | RNA Integrity Number |
| COD | Cause of Death |
| PMI | Post-mortem Interval |

**Extended Data Figure 1.**
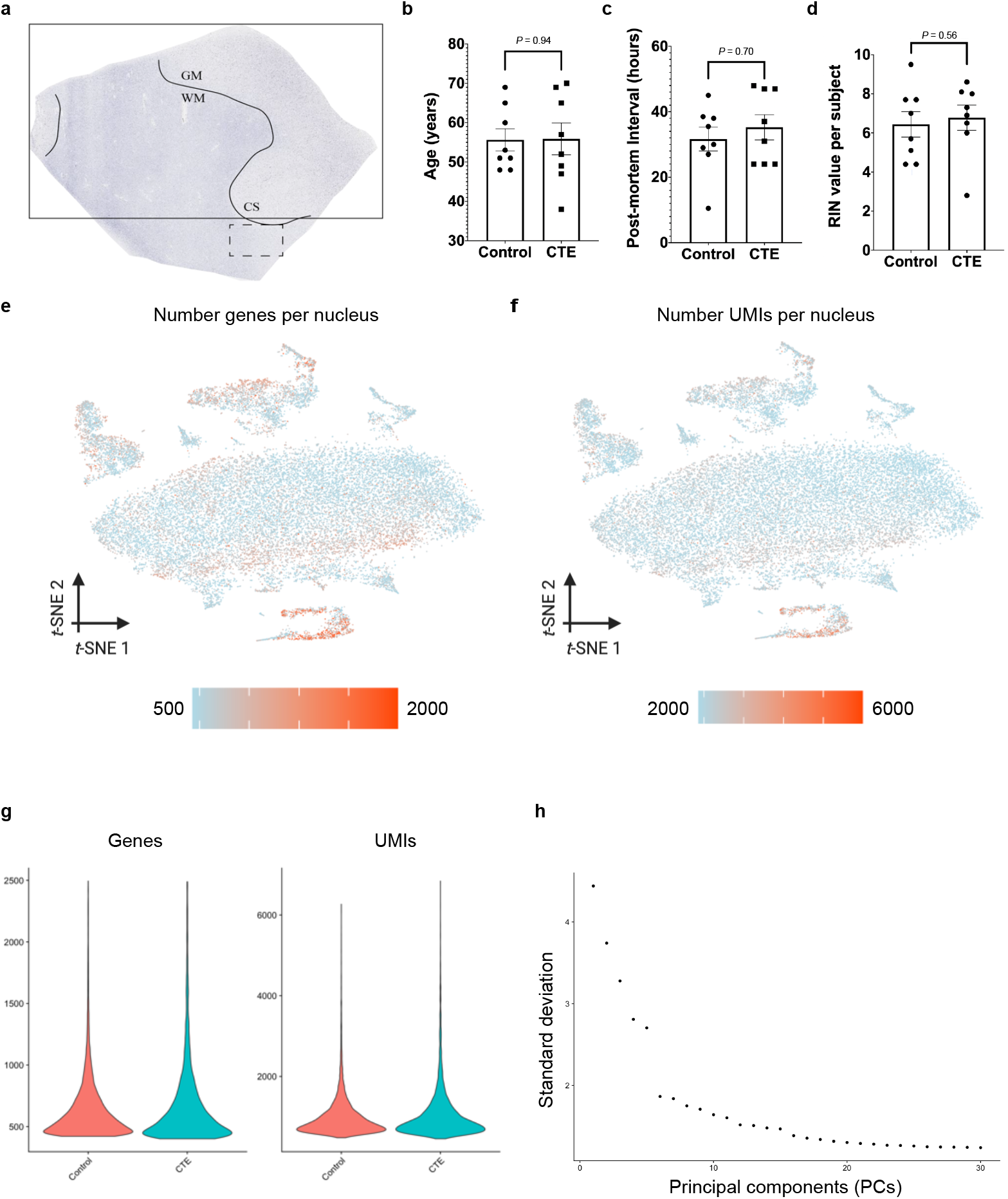
Tissue collection and quality control. **a.** Image of dorsolateral frontal white matter and depth of a cortical sulcus representing the region of tissue used for this project. Dashed rectangle represents a region taken for snRNA-seq analysis (only white matter at the depth of a cortical sulcus), solid rectangle represents a region taken for all in situ validation (gray and white matter adjacent to the region taken for snRNA-seq). GM: gray matter, WM: white matter, CS: cortical sulcus. **b.** Scatter plot with bar for each subject by age (*n* = 8 per condition, median of CTE = 54.5 years, median of control = 52.0 years, *P* = 0.94, two-tailed Mann-Whitney U test, *U* = 31.0). **c.** Scatter plot with bar for each subject by post-mortem interval (PMI) (*n* = 8 per condition, median of CTE = 34.0 hours, median of control = 32.75 hours, *P* = 0.70, two-tailed Mann-Whitney U test, *U* = 25.0). **d.** Scatter plot with bar for each subject by RNA integrity number (RIN) value (*n* = 8 per condition, median of CTE = 7.1, median of control = 6.4, *P* = 0.56, two-tailed Mann-Whitney U test, *U* = 26.0). **e.** tSNE projection colored by number of genes present in each nucleus. **f.** tSNE projection colored by number of unique molecular identifiers (UMIs) present in each nucleus. **g. (left)** Violin plots for the number of genes expressed in each nucleus for control and CTE subjects. **(right)** Violin plots for the number of UMIs present in each nucleus for control and CTE subjects. **h.** Elbow plot for the amount of variance produced by each principal component. Principal components (PCs) from 1 to 30 on the x-axis and standard deviation on the y-axis. All data was presented in Mean ± S.E.M.

**Extended Data Figure 2.**
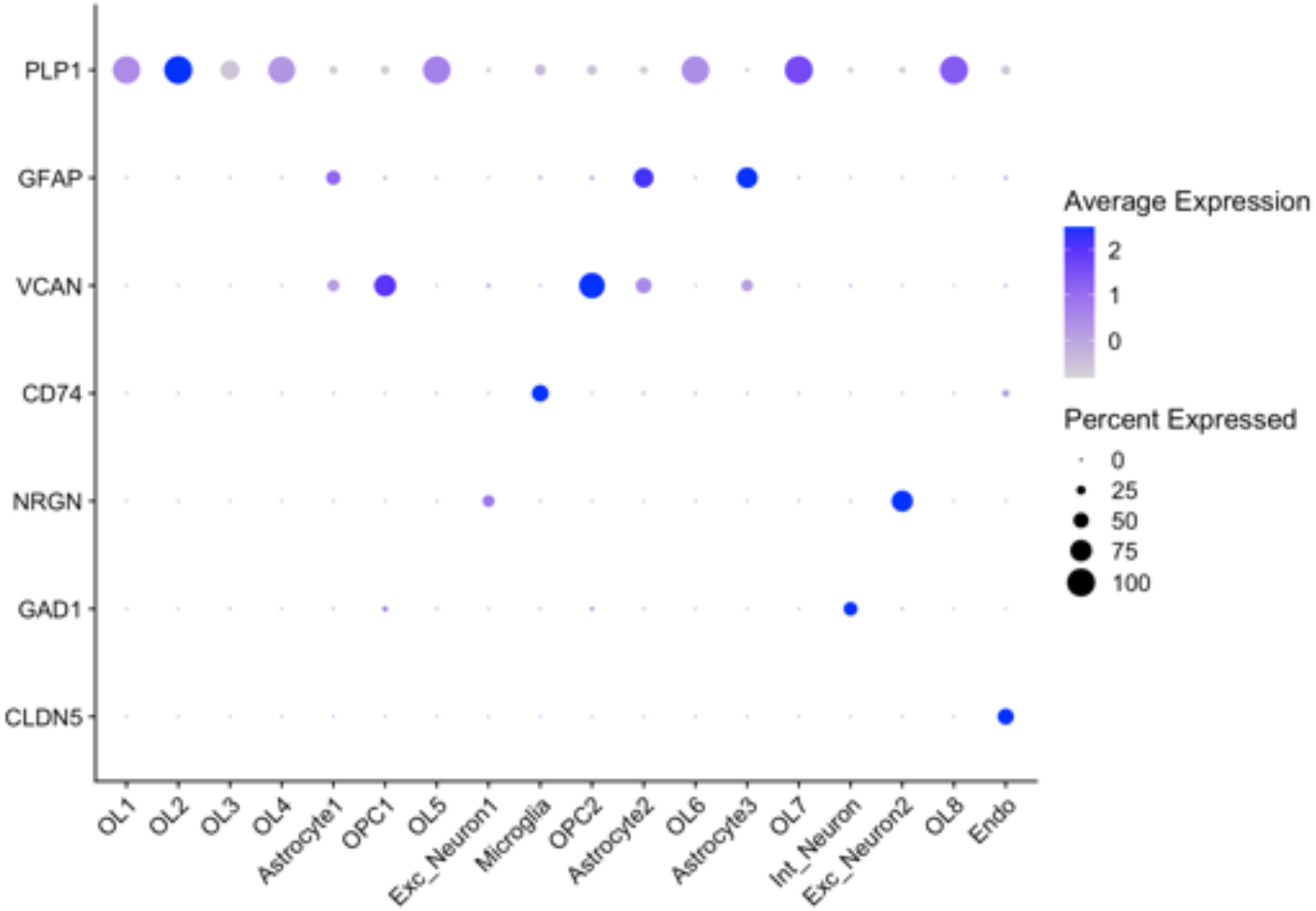
Cell-type-specific marker gene expression. Dotplot with cell types on the x-axis and marker genes on the y-axis. Size of dot relates to the percent of nuclei in a given cluster that express the corresponding marker gene. Color of dot relates to the average marker gene expression for the nuclei expressing the marker gene in a given cluster

**Extended Data Figure 3.**
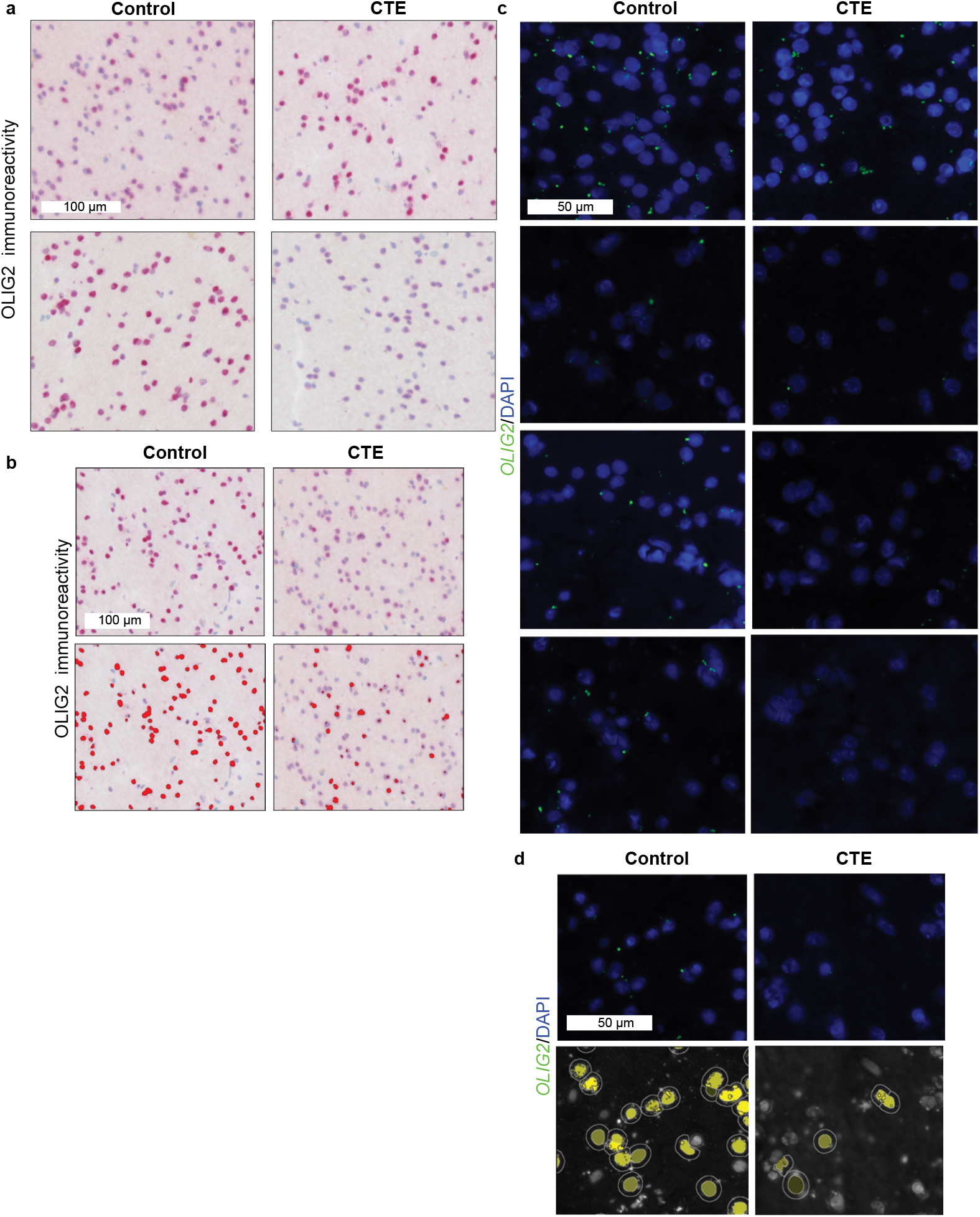
Validation of identification of fewer oligodendrocytes in CTE white matter compared to control. **a.** Representative immunohistochemistry images for control and CTE stained with anti-OLIG2 for oligodendrocyte lineage cells. Scale bar **(top-left, white)**: 100 μm. **b.** Immunohistochemistry images from main text **(top)** and the same images with masks **(bottom)** to demonstrate how positive cells are identified. Scale bar **(top-left, white)**: 100 μm. **c.** Representative smFISH images for *OLIG2* (green) and DAPI (blue). Scale bar **(top-left, white)**: 50 μm. **d.** smFISH images from main text **(top)** and the same images with masks **(bottom)** to demonstrate how nuclei and positivity are identified. Scale bar **(top-left, white)**: 50 μm.

**Extended Data Figure 4.**
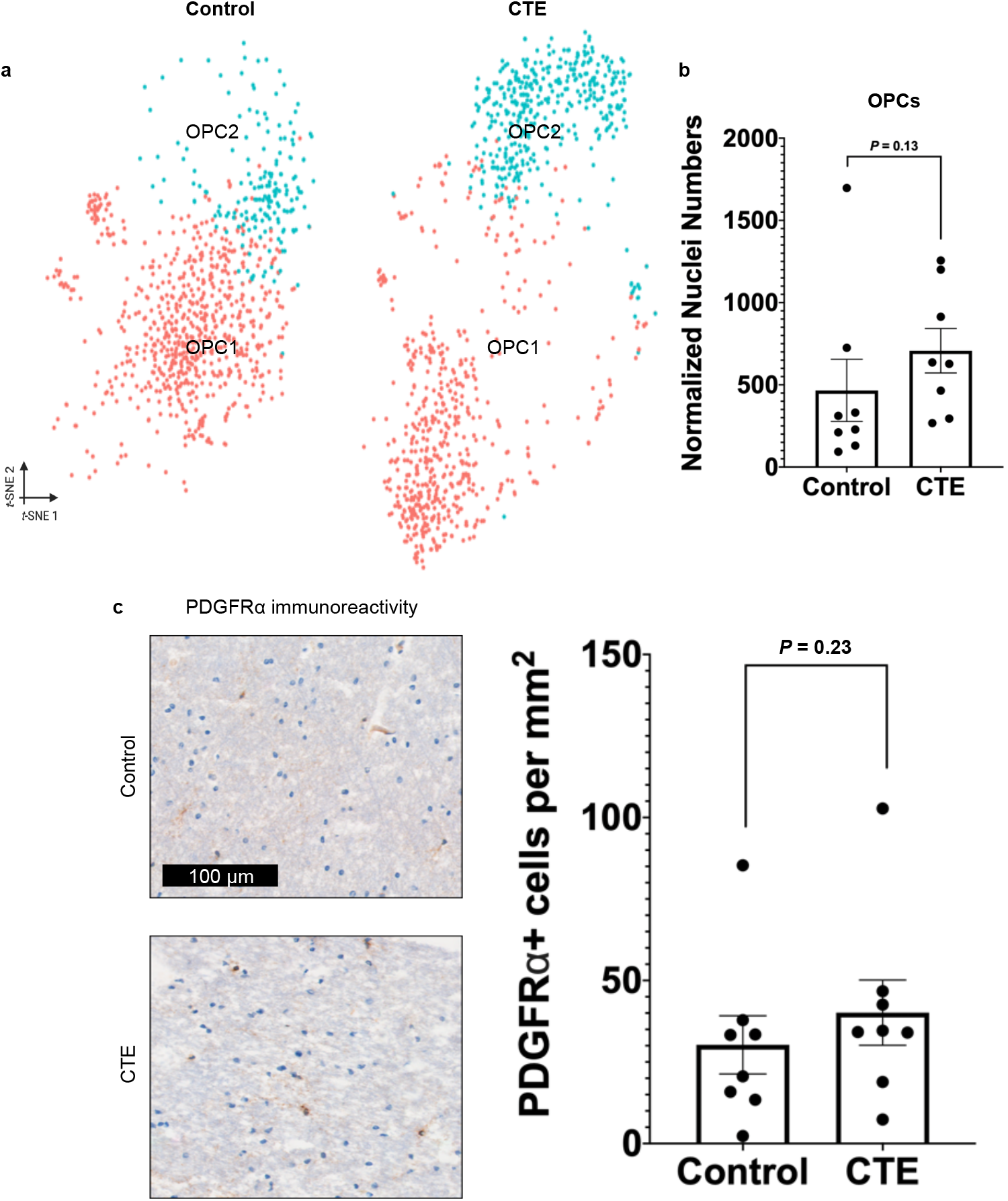
Oligodendrocyte precursor cell numbers do not change between conditions. **a.** tSNE projection of OPC nuclei subset from primary tSNE from Fig. 1b. **b.** Scatter plot with bar for normalized nuclei counts for all OPC nuclei, each dot represents the total normalized number of OPC nuclei in one subject (*n* = 8 per condition, *P* = 0.13, FDR-corrected two-tailed Mann-Whitney U test, *U* = 17.0). **c.** Representative immunohistochemistry images for control and CTE stained with anti-PDGFRα for OPC. Scale bar (**top, black)**: 100 μm. Quantification **(right)** for the number of PDGFRα-positive cells per mm^2^ over the entire white matter sample for each subject (*n* = 8 per condition, *P* = 0.24, one-tailed Student’s t-test, t(14) = 0.7363). All data was presented in Mean ± S.E.M.

**Extended Data Figure 5.**
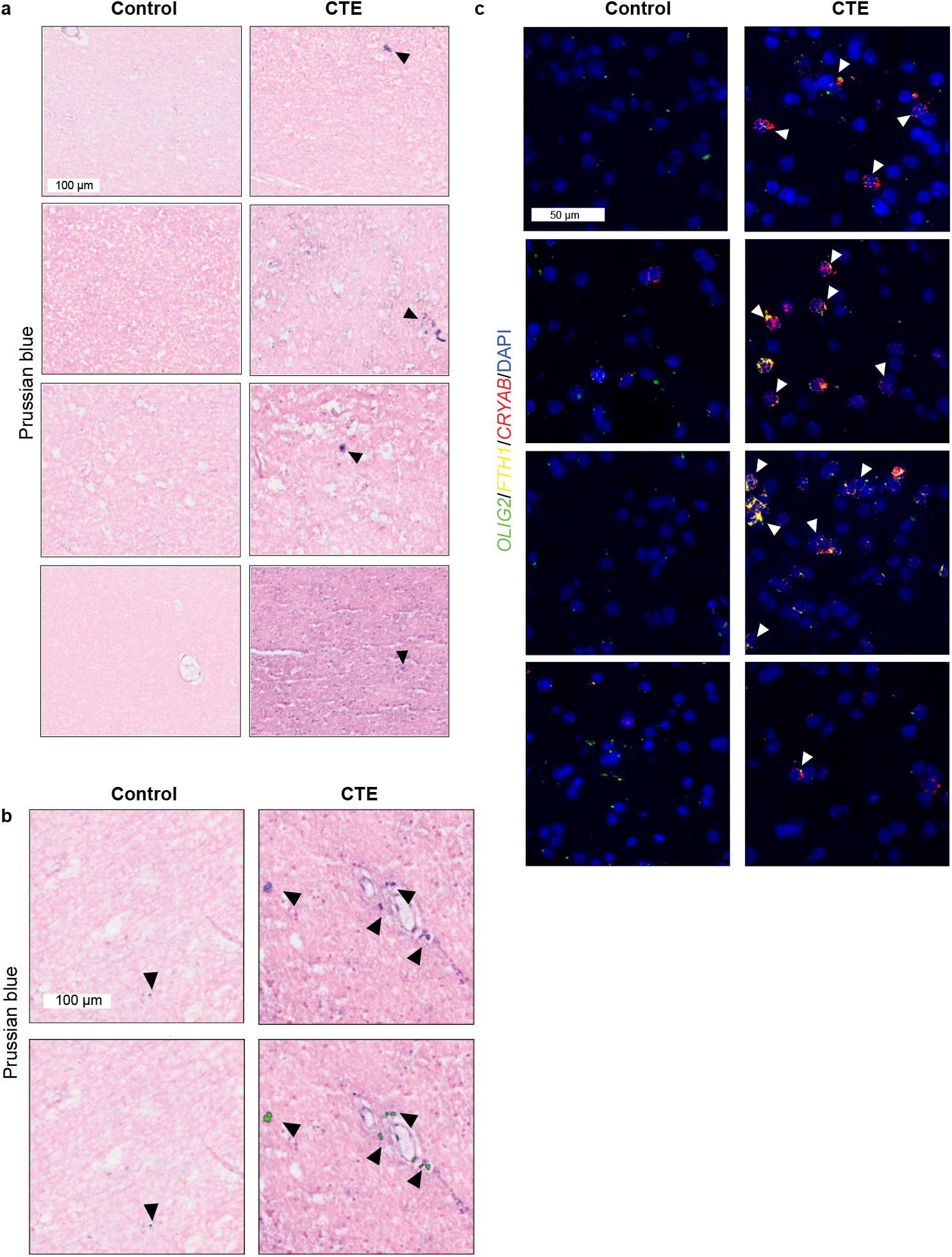
Validation of iron accumulation in CTE-specific oligodendrocytes. **a.** Representative images of Prussian blue staining for control and CTE. Black arrows label iron inclusions. Scale bar **(top-left, white)**: 100 μm. **b.** Prussian blue images from main text **(top)** and the same images with masks to demonstrate how iron inclusions are identified **(bottom)**. Scale bar **(top-left, white)**: 100 μm. **c.** Representative multiplexed smFISH images for *OLIG2* (green), *FTH1* (yellow), *CRYAB* (red), and DAPI (blue). White arrows label nuclei positive for *OLIG2*, *FTH1*, *CRYAB*, and DAPI. Scale bar **(top-left, white)**: 50 μm.

**Extended Data Figure 6.**
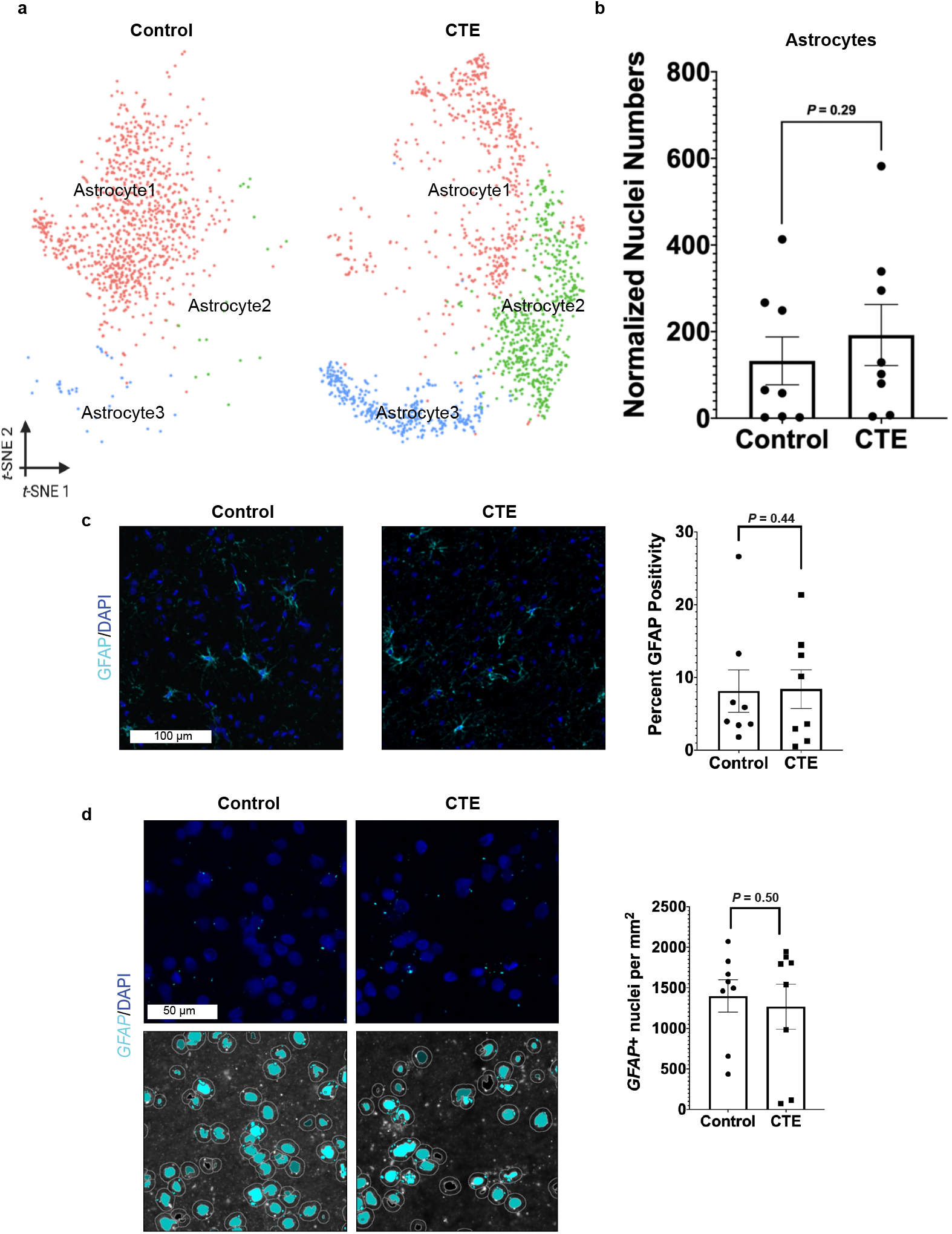
Astrocyte cell numbers do not change between conditions. **a.** tSNE projection of astrocyte nuclei subset from primary tSNE from Fig. 1b. **b.** Scatter plot with a bar for normalized nuclei counts for all astrocytes, each dot represents the total normalized number of astrocyte nuclei in one subject (*n* = 8 per condition, *P* = 0.29, FDR-corrected two-tailed Mann-Whitney U test, *U* = 0.29). **c.** Representative IF images for GFAP (teal) and DAPI (blue) in control and CTE. Scale bar **(left)**: 100 μm. Quantification **(right)** for percent of total area positive for GFAP IF for each subject (*P* = 0.44, one-tailed Student’s t-test). **d. (top)** Representative smFISH images for *GFAP* (teal) and DAPI (blue). **(bottom)** Same images as top but with masks to demonstrate how nuclei and positivity are identified. Scale bar **(top-left, white)**: 50 μm. Quantification **(right)** for number of *GFAP*-positive nuclei for each subject (*n* = 8 per condition, median of CTE = 1,667, median of control = 1,536, *P* = 0.50, one-tailed Mann Whitney U-test, *U* = 32.0). All data was presented in Mean ± S.E.M.

**Extended Data Figure 7.**
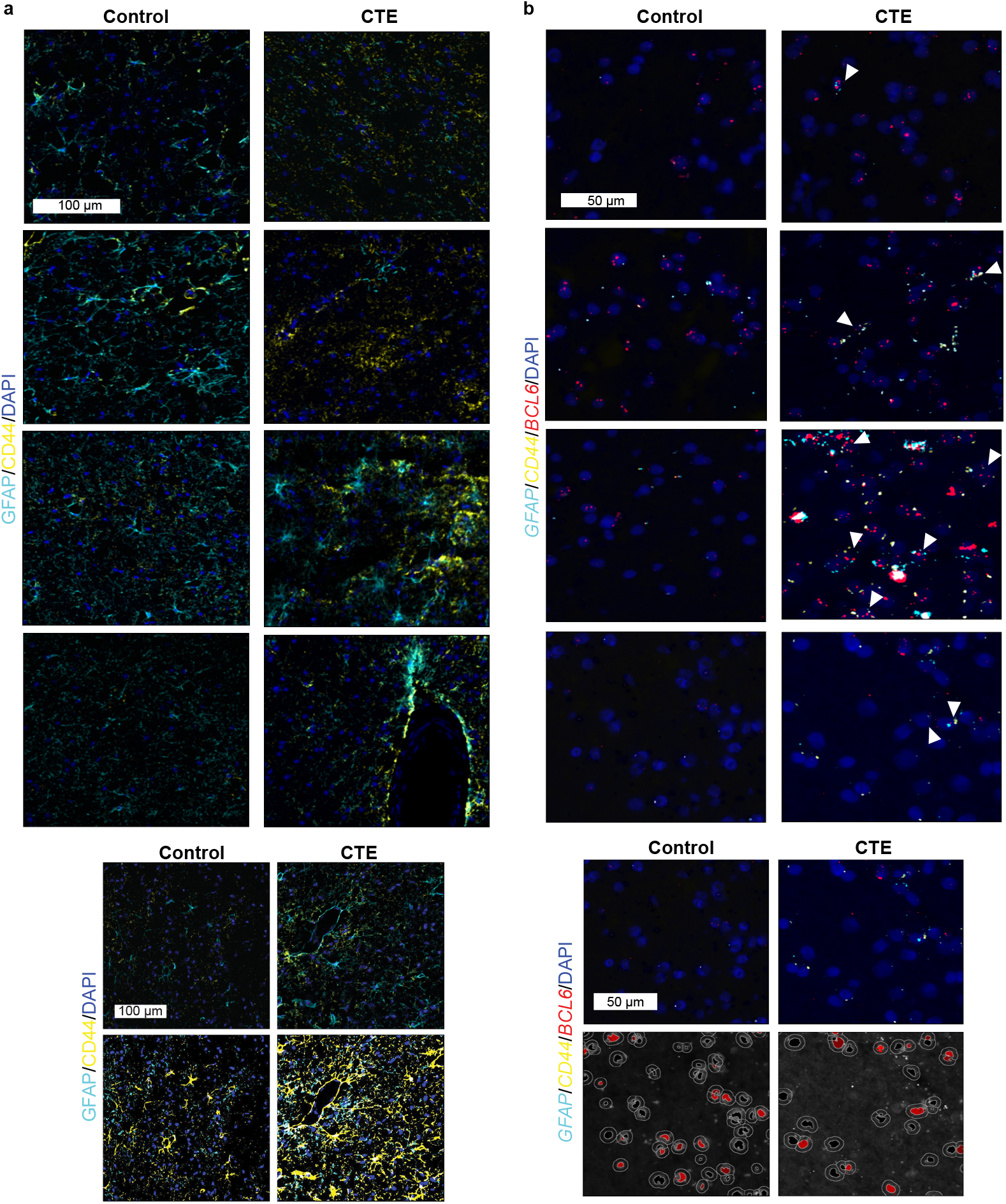
Validation of neuroinflammatory astrocytes in CTE white matter. **a.** Representative IF images for GFAP (teal), CD44 (yellow), and DAPI (blue) in control and CTE. Scale bar **(top-left, white)**: 100 μm. **b.** IF images from main text **(top)** and the same images with masks **(bottom)** to demonstrate how GFAP and CD44 positivity are identified. Scale bar **(top-left, white)**: 100 μm. **c.** Representative multiplexed smFISH images for *GFAP* (teal), *CD44* (yellow), *BCL6* (red), and DAPI (blue) in control and CTE. White arrows label nuclei positive for *GFAP*, *CD44*, *BCL6*, and DAPI. Scale bar **(top-left, white)**: 50 μm. **d.** Multiplexed smFISH images for *GFAP* (teal), *CD44* (yellow), *BCL6* (red), and DAPI (blue) in control and CTE **(top)** and the same images with masks **(bottom)** to demonstrate how nuclei and *BCL6* positivity are identified. Scale bar **(top-left, white)**: 50 μm.

## SUPPLEMENTARY INFORMATION LEGENDS

**Supplementary Table 1**: **snRNA-seq quality control**

**Supplementary Table 2**: **Cell-type marker genes**

**Supplementary Table 3**: **Differential gene expression analysis**

**Supplementary Table 4**: **Gene ontology**

**Supplementary Table 5**: **Normalized nuclei counts**

**Supplementary Table 6**: **Control counts matrix**

**Supplementary Table 7**: **CTE counts matrix**

**Supplementary Table 8**: **In situ validation data**

